# The mouse oocyte Balbiani body expands in growing follicles and influences nuclear position

**DOI:** 10.64898/2026.09.02.748954

**Authors:** Qi Yin, Allan C. Spradling

**Affiliations:** Howard Hughes Medical Institute Research laboratories, Department of Embryology, Carnegie Institution for Science, Baltimore, MD 21218, USA

## Abstract

The Balbiani body (Bb), a conserved oocyte structure enriched in organelles, forms during zygotene but behaves differently in several animal groups. The mouse Bb arises normally, but becomes smaller, with less organelle diversity, as primordial follicles enter a quiescent stage compared to Bbs from active lower vertebrate oocytes. In the oocytes of primary follicles that have returned to growth, we show that the primordial Bb remains organized around the same microtubule-organizing center (MTOC), but now acquires diverse organelles whose quality the Bb is postulated to improve. Thus, the mouse oocyte Bb’s structure is influenced by the primordial follicle’s increasing quiescence, but after the follicle returns to growth, the Bb resembles the Bb in zebrafish and *Xenopus* oocytes. Eventually, the mature Bb disperses as its MTOC relocates from the perinuclear region to the oocyte cortex, driven by opposing microtubule and actin forces that also reposition the nucleus to the oocyte center. We suggest that the mature Bb persists in each growing oocyte until its function in mitochondrial and organelle quality control is complete.

## INTRODUCTION

Oocyte production requires extensive molecular and cellular activities, including epigenetic reprogramming of germ cell chromatin, regulated movement and translation of maternal mRNAs, and acquisition of high-quality organelles in large numbers. The Balbiani body (Bb) is a composite structure found in oocytes of most animals, from zygotene through much of the oocyte growth period. It uses a core of microtubules to move and regulate mRNAs, including germ plasm mRNAs, and organelles (reviewed in Kloc et al., 2004; Jamieson-Lucy and Mullins, 2019; Spradling et al., 2022; Lei et al., 2024; Kar and Elkouby, 2026).

Because oocytes in different species develop at different rates and vary in the level of Bb involvement in different tasks at different times, the Bb’s appearance sometimes differs between species and developmental stages. In zebrafish and *Xenopus*, the Bb initially forms in zygotene oocytes and matures in oocytes of early follicles in a process dependent on the *Bucky ball* or *Xvelo* genes, which are required for a stable Bb structure, normal oocyte patterning and fertility (Marlow and Mullins, 2008; Boke et al., 2016). While phase separation undoubtedly plays a role (Kar et al., 2025), the enormous size of the Bb and its dynamic changes in organelle, mRNA and germ granule content, indicate that other mechanisms are also important. However, these phases are united by the extensive involvement of microtubule-dependent processes, which, together with the actin cytoskeleton, move and regulate mRNA ribonucleoprotein granules and organelles such as ER, Golgi, and mitochondria.

The fundamental relationship between the Bb and microtubule-organizing centers (MTOCs) has been documented in multiple species. The Bb is first associated with the centrosomes that cluster meiotic chromosome telomeres, even in species that do not form germline cysts (Tworzydlo et al., 2014). In zebrafish, microtubules contribute to the initial assembly and organization of the Bb, while *Bucky ball* promotes its stability and function through molecular condensation (Marlow and Mullins, 2008; Elkouby et al., 2016; Kar et al., 2025). In *Drosophila*, the transfer and accumulation of mitochondria and other organelles into a Bb depend on microtubules (Cox and Spradling, 2003). The number of mitochondria in the Bb can be manipulated using the Milton/TRAK1 adaptor protein, which links mitochondria to both plus- and minus-end-directed microtubule motors (Cox and Spradling, 2006). In mouse, the Bb forms during perinatal oogenesis around centrosomes (Pepling et al., 2007; Kloc et al., 2008; Niu and Spradling, 2022; Lei et al., 2024). Microtubule-dependent transport is required for normal mouse Bb formation (Lei and Spradling, 2016).

Perhaps the most sophisticated and least understood of the Bb-associated processes is organelle rejuvenation. Mitochondria are a particular focus of rejuvenation, as they contain their own genetic material and accumulate mutations at a higher rate than nuclear DNA, necessitating mechanisms for mitochondrial quality control and selection during oogenesis. The Bb has long been thought to help maintain oocyte mitochondrial quality (Cox and Spradling, 2003; Kloc et al., 2004; Pepling et al., 2007; Tworzydlo et al., 2016; Lieber et al., 2019; Palozzi et al., 2022; Monteiro et al., 2023; Sekula et al., 2024). The involvement of the Bb in rejuvenation of other organelles has only been postulated (Cox and Spradling, 2003), but little is understood beyond the theoretical importance of resynthesizing organelle molecular components from small molecules.

In mice and many other animal groups, the nucleus in most dictyate oocytes occupies an asymmetric position above the Bb and closer to the future animal pole. Later, in growing mouse oocytes, nuclei lie either eccentrically near the surface, or centrally. Toward the end of oocyte growth, nuclear centralization is regulated by actin (Almonacid et al., 2015, 2019). However, the role of MTOCs, microtubules and actin in positioning the nuclei of growing mammalian oocytes remains poorly understood.

In this study, we identify an MTOC-centered, mature Bb in growing mouse oocytes, comprising mitochondria, Golgi apparatus, endoplasmic reticulum (ER), and endosomes. We identify a similar central MTOC in the primordial Bb of oocytes from primordial follicles, follow the evolution of Bb organization during quiescence, and argue that the Bb dynamics are enabled by stable acetylated microtubules organized by a stable MTOC. MTOC activity eventually declines, and the MTOC relocates to the oocyte cortex at various times in individual growing oocytes, after which organelles disperse throughout the cytoplasm. The changes in the oocyte cytoskeleton associated with dispersal cause the nucleus to take up a central location in the mouse oocyte, making its location a marker for Bb presence. Our findings show that the Bb persists in mouse oocytes for a much longer period than previously supposed, supporting the idea that this organelle functions in a similar manner in diverse animals.

## RESULTS

### The oocytes from growing follicles contain a mature Bb

We studied the organelle distribution and cytoskeleton of oocytes from primary and secondary follicles to look for the presence of a mature Bb resembling the one in zebrafish and *Xenopus* (Figure 1). A single focus of γ-tubulin is located near the oocyte nucleus and is surrounded by a substantial localized region near the nucleus enriched in F-actin, as indicated by phalloidin staining, and the Golgi apparatus, as indicated by GM130 (Figure 1A). The presence of mitochondria in the same zone was suggested by low-level DAPI staining and confirmed by TFAM and TOMM20 immunofluorescence (Figure 1B), as well as by MitoTracker and tetramethylrhodamine methyl ester (TMRM) staining (Figure S1A,B). ER enrichment was documented by Calnexin staining (Figure 1C). Electron microscopy further verified the presence of mitochondria and Golgi-associated structures (Figure 1D). Further characterization confirmed that the structure consists of a central MTOC closely surrounded by mitochondria, with the Golgi apparatus and endosomes forming the outer layers (Figure 1E). The structure subsequently dispersed as oocytes developed (Figure S1C). We conclude that an MTOC-centered Bb resembling the one in zebrafish and *Xenopus* is found in growing mouse oocytes. Very recently, another group reported observing a similar structure (Zollo et al., 2026).

**Figure 1.**
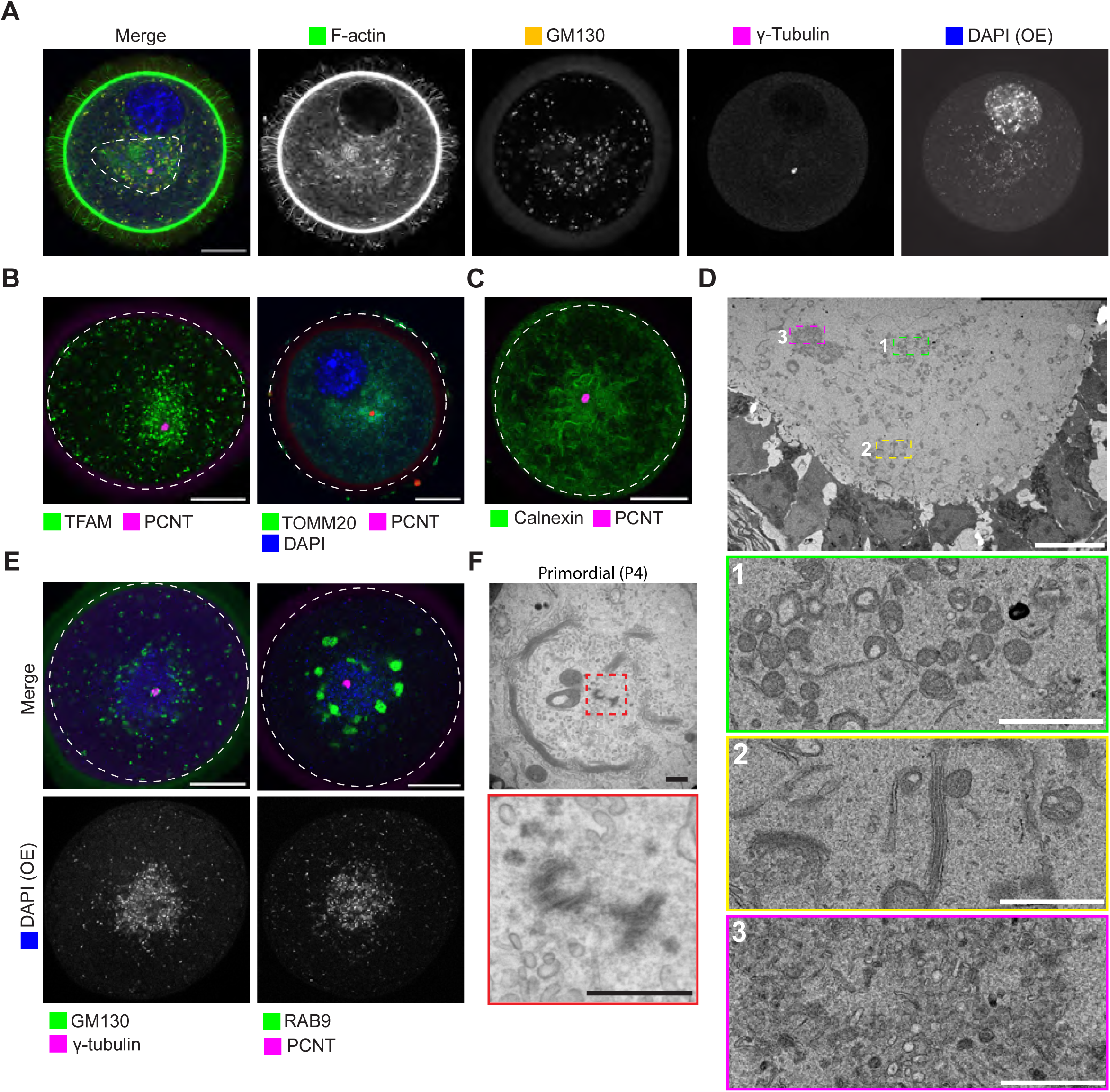
Characterization of the mature Balbiani body (Bb) in growing oocytes from secondary follicles. (A) Representative images showing the MTOC surrounded by mitochondria, the Golgi apparatus, and F-actin. Dashed line outlines the Bb. DAPI (OE), high laser power and long exposure. Scale bar, 20 μm. (B) Immunostaining for the mitochondrial markers TFAM and TOMM20. Dashed lines outline the oocytes. Scale bars, 20 μm. (C) Immunostaining for the endoplasmic reticulum (ER) marker Calnexin showing its presence within the mature Bb. Dashed lines outline the oocytes. Scale bar, 20 μm. (D) Representative electron micrographs showing the ultrastructure of the mature Bb in oocytes of secondary follicles, including mitochondria (enlarged in 1), the Golgi apparatus (enlarged in 2), and a multivesicular cluster (enlarged in 3). Scale bar, 10 μm; insets, 2 μm. (E) Representative images of the Golgi apparatus marker GM130 and the endosome marker RAB9, showing the spatial organization of the Golgi apparatus and endosomes relative to mitochondria and the MTOC. Dashed lines outline the oocytes. Scale bars, 20 μm. (F) Electron micrograph of a primordial Bb containing Golgi rings centered on a centriole pair (enlarged in the lower panel) in an oocyte from a primordial follicle of a P4 mouse. Scale bar, 500 nm.

A Bb had previously been identified in oocytes of primordial follicles (Figure 1F). This raises the question of how these two Bbs are related.

We studied the persistence of the Bb in oocytes of primordial follicles (Figure S2) by first examining its distinctive ring-shaped Golgi apparatus and centrioles (Pepling et al., 2007; Kloc et al., 2008). We carried out a series of EM and light microscopic studies on the oocytes of primordial follicles during the first four months of postnatal life to address Bb stability. Initially, in P4–7 (postnatal) oocytes, the Golgi elements associated with the Bb showed some variation in structure (Figure S2A). The rings, actually oblate spheroids, contained 4–5 pairs of Golgi stacks in cross section that were usually associated loosely along their edges; occasionally a pair of stacks was missing or was off to one side. Mitochondria observed in the EM sections varied widely in distribution (Figure S2A). The extent to which organelles are included in the image plane is summarized in three categories in the bar graph at the right, based on the assumption that the EM sections represent a random sample of planes containing the Golgi ring (Figure S2A, right). The Golgi rings persisted in oocytes of primordial follicles with relatively little change for at least 2 weeks. The organization of the stacks and their GM130 staining level diminished during the next 4 months but the stacks remained recognizable (Figure S2B,C). Two centrioles were identified in a perinuclear location, surrounded by the Golgi apparatus in oocytes of primordial follicles at three time points investigated by EM--P4, P7, and P19 (Figures 1F and S2D). We term the Bb in oocytes of primordial follicles a “primordial Bb” because it contains fewer organelles than the mature one in growing oocytes.

### The Bb expands using the central MTOC of the primordial Bb in growing oocytes

To gain more insight into the relationship between the primordial Bb seen in oocytes of primordial follicles and the mature Bb in oocytes of growing follicles, we examined changes in microtubule organization during this transition (Figure 2). Microtubules were sparse in oocytes of primordial follicles but increased substantially following follicle activation, with α-tubulin staining indicating microtubule enrichment on one side of the oocytes (Figures 2A, S3, and S4). Dense microtubule bundles were frequently associated with invaginations of the nuclear envelope (Figure 2B), resembling the nuclear envelope deformations associated with nuclear movement in *Drosophila* oocytes (Zhao et al., 2012). In contrast, in oocytes of secondary follicles whose nucleus had moved to the center, microtubules were evenly distributed throughout the ooplasm (Figure 2A, right).

**Figure 2.**
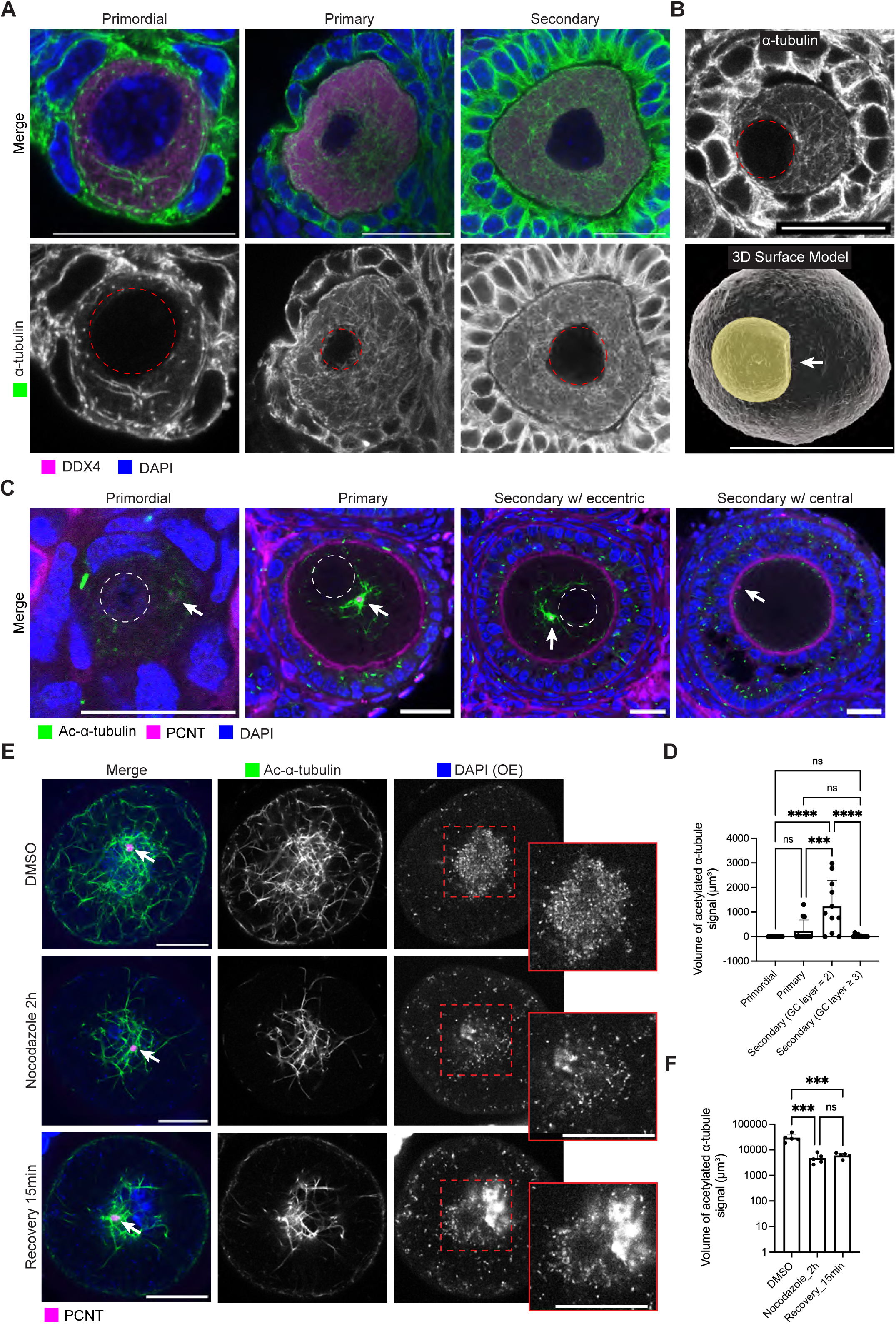
Microtubule organization and expansion of the primordial Bb from its single MTOC. (A) Representative images showing microtubule organization during follicle development. Dashed lines outline the nuclei. Scale bars, 20 μm. (B) Three-dimensional reconstruction of a primary follicle oocyte showing a nuclear indentation. Top, single optical section; bottom, 3D surface rendering generated using Imaris. Dashed lines outline the nucleus. Yellow shading indicates the nucleus. Arrow indicates the nuclear indentation. Scale bar, 20 μm. (C) Representative images of acetylated α-tubulin during follicle development. Dashed lines outline the nuclei. Arrows indicate MTOCs. Scale bars, 20 μm. Primordial, primordial follicle; Primary, primary follicle; Secondary, secondary follicle; Secondary w/ eccentric, secondary follicle containing an oocyte with an eccentric nucleus; Secondary w/ central, secondary follicle containing an oocyte with a centrally located nucleus. (D) Statistical analysis of acetylated α-tubulin around MTOCs in oocytes during follicle development. Primordial, primordial follicles, n = 14; Primary, primary follicles, n = 13; Secondary (GC layer = 2), secondary follicles with 2 layers of granulosa cells (GCs), n = 11; Secondary GC (GC layer ≥ 3), secondary follicles with 3 or more layers of granulosa cells, n = 10. *P* values were determined using one-way ANOVA, followed by Tukey’s multiple comparisons test. (E) Representative images of oocytes from secondary follicles following 2 h nocodazole treatment and 15 min of recovery. DAPI (OE), high laser power and long exposure. Arrows indicate MTOCs. Scale bars, 20 μm. (F) Statistical analysis of total acetylated α-tubulin of oocytes from secondary follicles treated with DMSO, nocodazole, or nocodazole followed by washout and 15 min recovery. n = 5 per group. P values were determined using one-way ANOVA, followed by Tukey’s multiple comparisons test. ns, not significant. *** *P* < 0.001, **** *P* < 0.0001.

We looked for a precursor to the MTOC of the growing oocyte in follicles stained for acetylated microtubules during the transition from primordial to growing follicles (Figure 2C). A single MTOC located at the center of the microtubule-enriched region was clearly visible in the oocytes of primordial follicles. Following follicle activation, acetylated α-tubulin (Lys40) staining in oocytes became prominent and was highly enriched at an MTOC (Figures 2C,D, and S3). Acetylated α-tubulin slowly decreased as oocytes progressed toward later stages. Thus, the MTOC of the primordial Bb becomes more active in organelle recruitment to produce the mature Bb.

We further examined the organization of microtubules and the Golgi apparatus in follicles at different stages of development (Figure S4). In the oocytes of primordial follicles, microtubules appeared to project outward from the center of the region surrounded by the Golgi stacks (Figure S4, “Primordial”). Shortly after activation, the Golgi stacks begin to break up, and the microtubules grow and extend outside the region (Figure S4, “Transitional”). In the oocytes of primary follicles, the Golgi has become further reduced in size, while abundant microtubules reside at the center of the oocyte, near an indentation in the nuclear membrane (Figure S4, “Primary”). The Golgi has dispersed further in secondary follicles, while the microtubule organization continues to correlate with nuclear asymmetry (Figure S4, “Secondary”). These studies strongly argue that the Bb in oocytes of growing follicles expands on a microtubule array stabilized by acetylation and centered on the MTOC of the oocytes of primordial follicles that was used to cluster the Golgi apparatus into a ring shape and to generate the primordial Bb.

We also tested whether microtubules are required to maintain the organization of the mature Bb in the oocytes of early secondary follicles (Figures 2E and S5A,C). Treatment with nocodazole substantially disrupted the microtubule network, but residual acetylated α-tubulin remained detectable (Figure 2E). Mitochondria dispersed following nocodazole treatment (Figures 2E and S5C), whereas removal of nocodazole led to partial re-enrichment of mitochondria around the MTOC (Figure 2E). Following nocodazole washout, microtubules regrew around the MTOC (Figure S5D,E), consistent with the ability of the MTOC to nucleate microtubules (Łuksza et al., 2013). Thus, disruption and subsequent recovery of microtubules were accompanied by corresponding dispersal and expansion of the MTOC-associated mitochondrial cluster.

Because F-actin was also enriched within the Bb (Figure 1A), we next asked whether actin contributes to the organization of the mitochondrial cluster of Bb in oocytes of secondary follicles. Unexpectedly, treatment with cytochalasin D did not disperse the mitochondrial cluster; instead, mitochondrial enrichment increased (Figure S5B,C). Thus, microtubule and F-actin perturbations produced opposite effects on mitochondrial organization, suggesting that these cytoskeletal systems exert opposing influences on the maintenance of the MTOC-associated Bb.

### Progressive nuclear centralization and MTOC relocation

Previous studies have reported that oocyte nuclei remain predominantly eccentric before the antral stage (Brunet and Maro, 2007). We therefore asked whether microtubules, MTOC, or the Bb contribute to nuclear positioning in oocytes during follicle development. Using whole-mount staining (Li et al., 2017), we examined oocyte nuclear positioning throughout follicle development (Figure 3A). In the oocytes of primary follicles, 19.4% of nuclei were centrally positioned, increasing to 38.6% in the oocytes of secondary follicles and 100% in the oocytes of antral follicles (Figure 3B,C).

**Figure 3.**
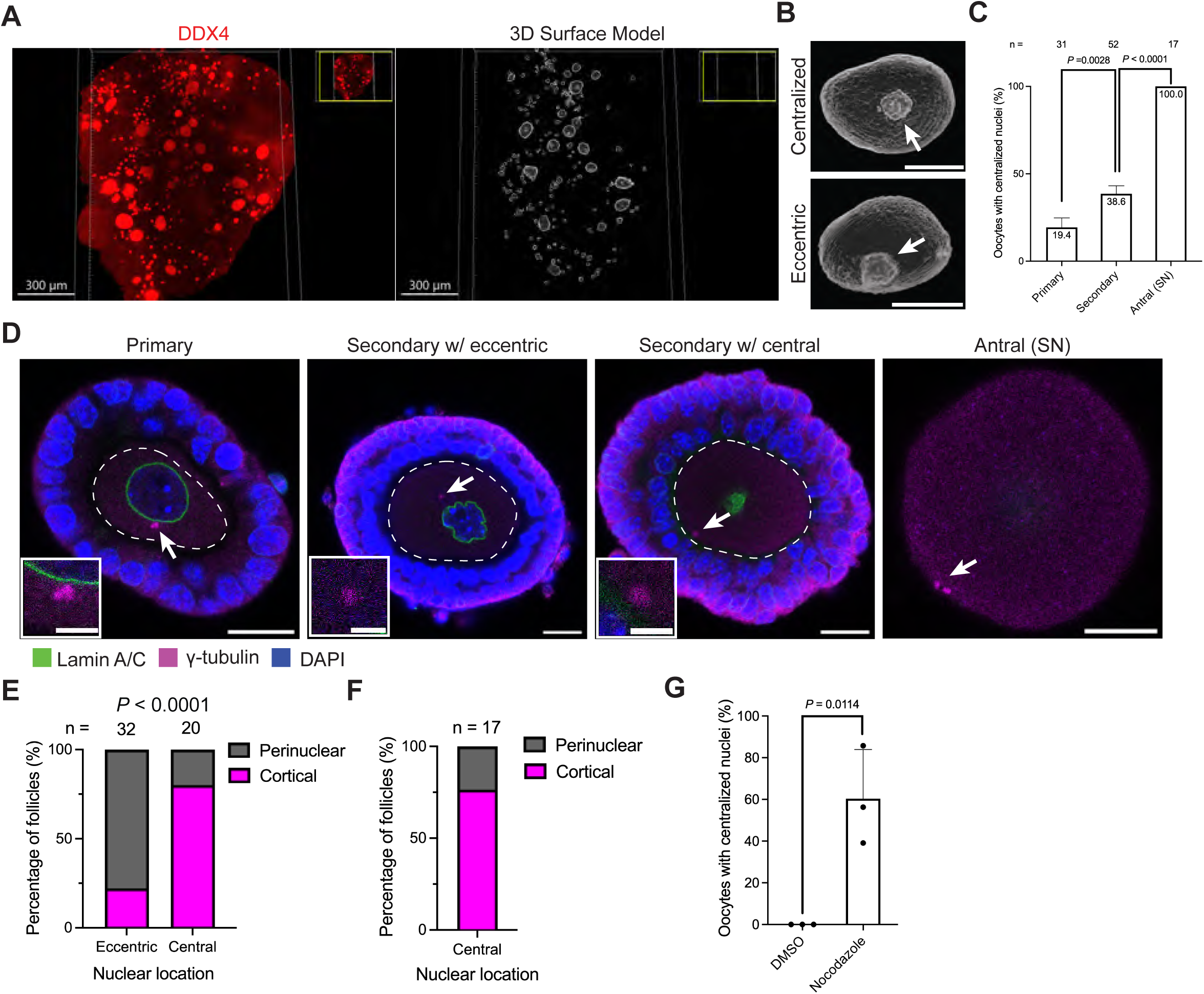
Nuclear centralization and MTOC redistribution during mouse follicle development. (A) Whole-mount immunostaining of the ovary (left) and corresponding 3D surface reconstruction (right). Scale bars, 300 μm. (B) Representative 3D surface reconstruction of oocytes generated using Imaris. Arrows indicate the nuclei. Scale bars, 5 μm. (C) Percentage of oocytes with centralized nuclei at different stages of follicle development. Numbers above the bars indicate the number of oocytes analyzed. Data were obtained from three mice. (D) Representative images of MTOCs and the nuclear envelope in follicles and oocytes. Dashed lines outline the oocytes. Arrows indicate MTOCs, enlarged in the insets. Scale bars, 20 μm; insets, 5 μm. (E) Quantification of MTOC localization in secondary follicles. Eccentric: Oocytes with eccentric nuclei; Central: Oocytes with centrally localized nuclei. Numbers above the bars indicate the number of oocytes analyzed. *P* value was determined using the chi-square test. (F) Quantification of MTOC localization in SN oocytes. The number above the bars indicates the number of oocytes analyzed. (G) Quantification of nuclear positioning in oocytes from secondary follicles following treatment with DMSO or 5 μM nocodazole for 2 h. Data were collected from three independent experiments. *P* values were determined using an unpaired, two-tailed Student’s *t* test. * *P* < 0.05, ** *P* < 0.01, *** *P* < 0.001. Primary, primary follicle; Secondary, secondary follicle; Secondary w/ eccentric, secondary follicle containing an oocyte with an eccentric nucleus; Secondary w/ central, secondary follicle containing an oocyte with a centrally located nucleus. Antral (SN), SN oocyte.

In oocytes of primary follicles, MTOCs were detected adjacent to the nucleus (Figure 3D). As follicles progressed to the secondary stage, MTOCs in some oocytes relocated to the oocyte cortex (Figures 3D and S6A). Comparison of MTOC and nuclear positioning revealed that MTOCs were predominantly perinuclear in oocytes with eccentric nuclei, whereas cortical MTOC positioning was more frequent in oocytes with centrally located nuclei (Figure 3E). In antral follicles, MTOCs predominantly localized to the cortex (Figure 3F).

We next investigated whether microtubules contribute to the maintenance of nuclear eccentricity in growing oocytes. After 2 h of nocodazole treatment, nuclei that were initially eccentric moved toward the center of the oocyte, suggesting that MTOCs and MTOC-associated microtubules contribute to maintaining nuclear eccentricity (Figure 3G). Overnight nocodazole treatment during in vitro follicle culture similarly resulted in nuclear centralization in oocytes (Figure S6B,C). Notably, MTOCs concomitantly relocated toward the cortex, while few acetylated microtubules remained around the MTOCs (Figure S6B,D), suggesting that MTOC-associated microtubules normally maintain MTOCs in the perinuclear region and thereby contribute to maintaining nuclear eccentricity.

### Composition of MTOCs changes during oocyte growth

During follicle growth, MTOCs in oocytes undergo changes in size and function, highlighting the importance of investigating their molecular composition (Figure 4A). We observed an increase in MTOC size during oocyte growth based on PCNT staining (Figure 4B). Most proteins previously identified in SN (surrounded nucleolus) oocytes (Gueth-Hallonet et al., 1993; Ge et al., 2010; Baumann et al., 2017; Bury et al., 2017; Wu et al., 2022; So et al., 2019; Gu et al., 2025) were also detected in MTOCs of NSN (non-surrounded nucleolus) oocytes (Figures 4C and S7). However, Ninein and PCM1 behaved differently (Figure 4D,E). Ninein, a microtubule-anchoring protein (Bouckson-Castaing et al., 1996; Lecland et al., 2019), remained localized to MTOCs through the NSN stage but was absent from MTOCs of SN oocytes, suggesting its involvement in maintaining microtubule-organizing activity during oocyte growth (Figure 4D). PCM1 has been shown to participate in the recruitment of proteins to centrosomes, including Ninein (Dammermann and Merdes, 2002). Consistent with this role, PCM1 was progressively lost from MTOCs during oocyte growth (Figure 4E), suggesting that compositional remodeling of MTOCs accompanies changes in the ability of MTOC to nucleate microtubules. Further studies are warranted to determine the specific roles of Ninein and PCM1 in regulating MTOC function during oocyte growth.

**Figure 4.**
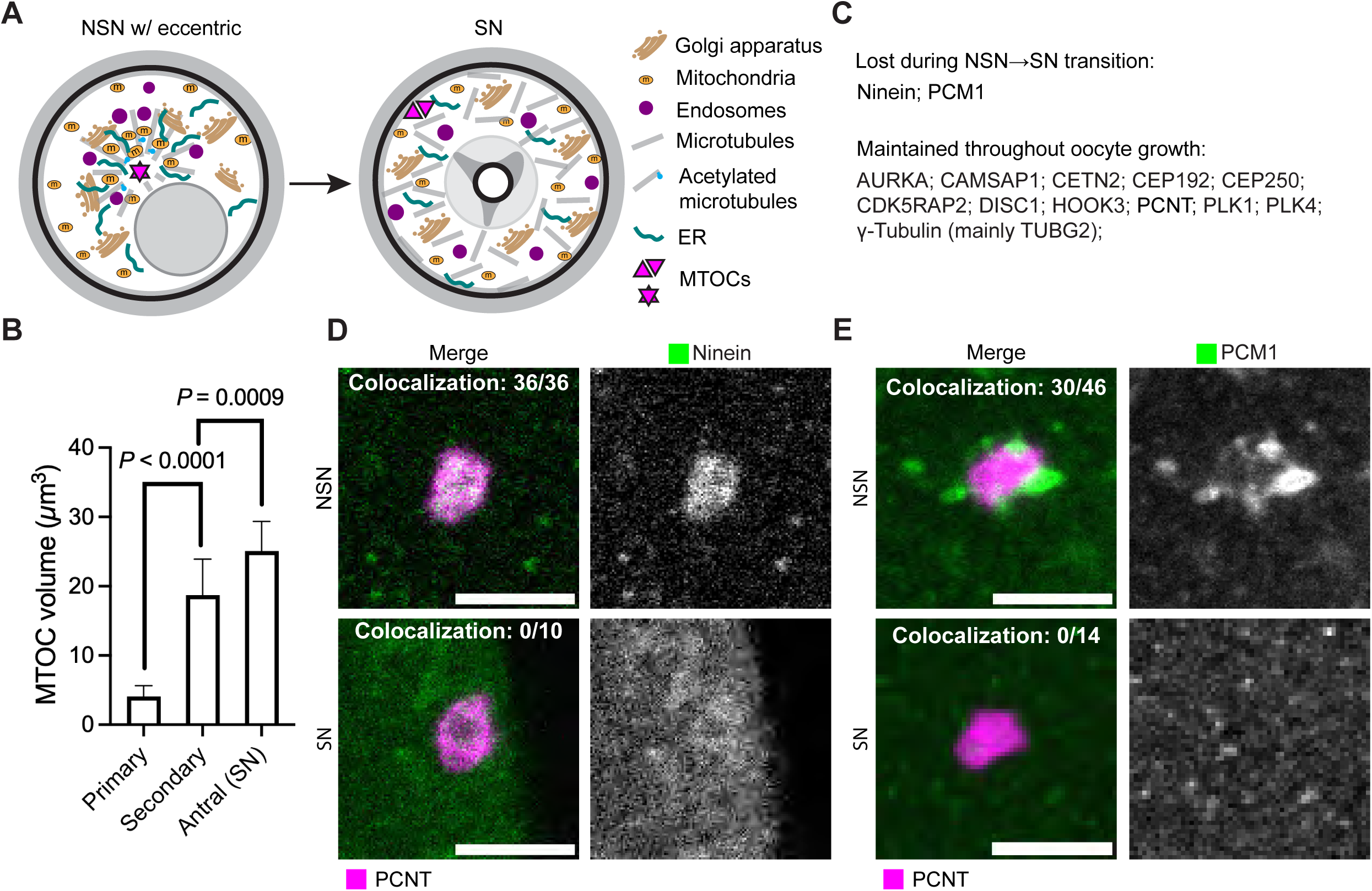
Changes in MTOC composition in oocytes during development. (A) Schematic summarizing the dynamics of the nucleus, organelles, and cytoskeleton during the transition from NSN to SN oocytes. In growing NSN oocytes, an MTOC hub forms in the center of the oocyte while the nucleus remains eccentric. MTOCs are surrounded by mitochondria, the endoplasmic reticulum, the Golgi apparatus, and endosomes. In SN oocytes, the nucleus becomes centrally positioned, whereas most MTOCs relocate to the cortex and the MTOC hub disperses. NSN w/ eccentric, NSN oocyte with an eccentric nucleus. (B) Quantification of MTOC volume during follicle development using Imaris based on pericentrin staining. Primary, primary follicles, n = 20; Secondary, secondary follicles, n = 27; Antral (SN), SN oocytes, n = 8. *P* values were determined using one-way ANOVA, followed by Tukey’s multiple comparisons test. (C) Summary of the dynamic changes in MTOC-associated proteins during follicle development. (D) Representative images showing the localization of Ninein in NSN and SN oocytes. Colocalization of Ninein and PCNT is shown in the images. Scale bar, 20 μm. (E) Representative images of PCM1 immunostaining in NSN and SN oocytes. Colocalization of PCM1 and PCNT is shown in the images. Scale bar, 20 μm. *** *P* < 0.001, **** *P* < 0.0001.

## DISCUSSION

### Growing mouse oocytes expand a pre-existing primordial Bb

We identified a common MTOC-centered Bb in primordial and growing mouse oocytes, that show major differences in appearance as oocytes pass through quiescence and enter rapid growth (summary: Table 1). Our findings suggest that the mouse Bb is dynamic, but fundamentally similar to the Bbs of other investigated vertebrates and many invertebrates, when compared at similar developmental stages of oogenesis and physiological environmental states. For example the striking prominence of the Golgi in primordial Bb but not in mature Bb, may reflect a developmental difference related to the transition to the relative quiescence of primordial follicles. This transition may require high secretory activity to produce an extracellular matrix promoting quiescence by blocking many extracellular signals (Niu and Spradling, 2020).

**Table 1.** Organelle and cytoskeletal dynamics in oocytes during follicle development.

|  |  | Oocytes in primordial follicles | Oocytes in primary follicles | Oocytes in secondary follicles | Oocytes in antral follicles (SN stage) |
|---|---|---|---|---|---|
| Nuclear location |  | Eccentric | Mostly Eccentric | Gradually centralized | Centralized |
| Bb | Mitochondria | Variable | Begin to accumulate around MTOC | Enriched around MTOC | Dispersed |
|  | Golgi apparatus | Golgi ring around MTOC | Golgi ring partially dissociates into Golgi dots | Enriched around MTOC (as dots) | Dispersed (as dots) |
|  | ER | Absent from Bb | Begin to become enriched | Enriched around MTOC | Dispersed |
|  | MTOC | Beside the nucleus | Increases in size | Located at the center of Bb | Relocates to the oocyte cortex |
| Microtubules |  | Short microtubules extending from MTOC | Microtubule become longer | Densely distributed throughout the oocytes | Microtubules become shorter |
| Acetylated microtubules |  | Absent | Begin to emerge from minus end | Stabilize microtubules around MTOC | Few remains in the oocyte cortex |

Very recently, Zollo et al. (2026) reported a similar organelle-rich structure in growing oocytes, which they termed the Zollo body. Our study was conducted without knowledge of their work and converges with their study in identifying an MTOC-centered compartment enriched in mitochondria, Golgi apparatus, and ER during oocyte growth. Although they refer to this structure as the Zollo body, we present evidence that it represents a mature Bb derived from the primordial Bb in oocytes of primordial follicles, based on developmental changes in organelle abundance using a constant MTOC-centered organization. Many other parts of these papers address independent issues.

We identified several differences between the Bb in oocytes of primordial and growing follicles. A major change was a reduction in the amount of Golgi material. The Golgi ring itself changed very little for at least 2 weeks, and then slowly reduced in size but was still present in the oocytes of primordial follicles even at 4 months. This reduction correlated with a smaller associated microtubule array. However, the Bb and its organelles probably continue to function in the oocytes of primordial follicles at a reduced level. Major changes in Golgi content occurred during expansion of the Bb in growing oocytes, but some of the primordial Golgi remained present, at least initially.

The longer active period of the mature Bb is indicated by the fact that greater than 80% of primary and 60% of secondary oocytes had asymmetrically located nuclei indicative of a Bb (Figure 3B). Only very late, at the SN stage, was the nucleus always located centrally, without evidence of a Bb. The expansion of the Bb in growing oocytes and its persistence show that a mitochondria-rich Bb is present for a much larger fraction of oogenesis than previously supposed, one much closer to the long-lived but dynamic Bb characteristic of zebrafish and *Xenopus* (Marlow and Mullins, 2008; Boke et al., 2016).

### The mature Bb may continue the process of mitochondrial and organelle improvement

The organization of the mature Bb around the same MTOC and its similar structure suggest that it continues to participate in mitochondrial and organelle quality selection. This process is known to persist at least up to the transition from early to late secondary follicles (Xie et al., 2025). Healthy mitochondria may be transported toward MTOC-associated regions through dynein-dependent mechanisms. Extensive actin and ER signals within the Bb may also facilitate mitochondrial fission and expansion of the mitochondrial population. In *Drosophila*, there is evidence that mitophagy participates in mitochondrial improvement during oogenesis (Lieber et al., 2019; Palozzi et al., 2022; Monteiro et al., 2023). Detailed comparison of the qualitative and quantitative differences between the functions of the Bb in oocytes of primordial and growing follicles awaits an improved understanding and assays addressing specific functions and their associated rates.

Differences between species and between individual oocytes in the duration of Bb presence may reflect the amount of time required to transfer and rejuvenate the cytoplasmic components of the oocyte. In *Xenopus*, the ring canals separating nurse-like cells and the oocyte do not expand in diameter like they do in *Drosophila* (Davidian and Spradling, 2025). Gaps in the membranes separating germ cells within a cyst have not been observed in *Xenopus* as they are in mouse, allowing passage of entire organelles (Lei and Spradling, 2016). Endosomes loaded with ER and Golgi proteins are found in *Xenopus* ring canals. Consequently, *Xenopus* oocytes may require a long period of time to transfer and rejuvenate oocyte organelles, creating a need for the Bb to persist in growing oocytes (Davidian and Spradling, 2025). Likewise, differences between individual growing oocytes in the time that the Bb persists may be determined by a feedback system that signals the Bb to dissociate when organelle improvement has been completed.

### The MTOC of the mature Bb is likely derived from the meiotic centrosomes of the early mouse oocytes

The identification of a single MTOC that appears to catalyze the organization of both Bbs in oocytes of primordial and growing follicles raises the question of its origin. Previous studies concluded that mouse centrosomes do not persist past pachytene based on the loss of centrioles detectable in electron micrographs (Szollosi et al., 1972). However, more sensitive studies looking at specific centrosome proteins show that some centrosomes do persist far longer. Post pachytene oocytes not only accumulate pericentrin-rich centrosomes that disperse as part of primordial Bb formation and its Golgi ring in oocytes of primordial follicles. They also contain two much smaller γ-tubulin positive structures that were proposed to be persistent meiotic centrosomes (Lei and Spradling, 2016). Two very similar structures containing CETN2 doublets were identified (Simerly et al., 2018) that persist into the antral and meiotic metaphase stages, although they do not contribute in a conventional way to meiotic spindles. Thus, a persistent meiotic centrosome pair in each oocyte that survives quiescence represents a strong candidate for the MTOC of the mouse Bb.

### The mouse oocyte cytoskeleton remodels the oocyte cytoplasm during oogenesis

Unlike *Drosophila* and other species in which nuclear positioning determines the dorsoventral axis of the embryo, the significance of nuclear positioning in mammalian oocytes remains incompletely understood. Previous studies have reported a relationship between oocyte developmental competence and nuclear position (Bellone et al., 2009; Kincade et al., 2023). Our studies now reinforce the need to further investigate the cytoplasmic polarity associated with oocyte nuclear eccentricity. During this time, the MTOC, microtubules including acetylated microtubules, the Golgi apparatus, and associated organelles form a polarized intracellular system that may have significance for later development. For example, it might allow directional secretion, including the secretion of zona pellucida proteins (El-Mestrah et al., 2002). This organization may also contribute to patterned communication between oocytes and surrounding granulosa cells during follicle development. Further studies are required to determine whether polarized secretion or organelle organization influences subsequent oocyte competence and embryonic development.

### The mouse oocyte transitions between several states of cytoplasmic organization during oogenesis

Once their nucleation activity declines, MTOCs in NSN oocytes appear to become relatively inert structures that can be displaced by actin-dependent forces (Almonacid et al., 2015, 2019). In SN oocytes entering meiotic maturation, MTOCs again interact extensively with microtubules (Wickramasinghe and Albertini, 1992), resembling the transition between interphase and mitotic states in somatic cells. In *Drosophila*, loss of MTOC activity is preceded by the disappearance of pericentriolar material proteins (Pimenta-Marques et al., 2016). In mouse oocytes, although some proteins, including PCM1 and Ninein, were lost from MTOCs during follicle development, widespread loss of PCM proteins was not observed, as proteins like PCNT, γ-tubulin, and CETN2 remained associated with MTOCs. Further investigation will be required to understand the structural organization of oocyte MTOCs.

Together, these findings suggest that mammalian oocyte growth involves a coordinated transition between distinct cytoplasmic organizational states. An MTOC-centered mature Bb forms after follicle activation, when microtubules organize mitochondria and other organelles and contribute to maintaining nuclear eccentricity. As oocytes grow, cytoskeletal remodeling relocates the MTOC to the cortex, disperses the Bb, and permits nuclear centralization. Thus, rather than serving only as a static organelle aggregate, the Bb may represent a dynamic organizational state that couples cytoskeletal architecture with organelle remodeling during the prolonged growth phase of mammalian oocytes.

## MATERIALS AND METHODS

### Experimental mice

All animal experiments were performed in accordance with the guidelines of the Institutional Animal Care and Use Committee of the Carnegie Institution of Washington (CIW). The Office of Laboratory Animal Welfare (OLAW) assurance number is D16-00494. Mice were housed in specific pathogen-free (SPF) facilities under a 12-hour light/dark cycle. C57BL/6J mice (Strain #: 000664) were purchased from The Jackson Laboratory. Only female mice were used in this study.

### Mouse follicle and oocyte collection

Follicles at different developmental stages were collected as previously described (Jaffe et al., 2011). Briefly, ovaries were dissected and maintained in M2 medium (MilliporeSigma, MR-015-D). Follicles from the ovarian periphery were gently teased away using needles. Individual follicles were collected using hand-pulled glass capillaries and processed for subsequent experiments. Follicles were classified according to established morphological criteria. Primary follicles contained a single layer of cuboidal granulosa cells; secondary follicles contained two to four layers of granulosa cells; preantral follicles contained more than four layers of granulosa cells without an antral cavity; and antral follicles contained an antral cavity. Unless otherwise stated, all follicle and oocyte experiments were performed using 2–4-week-old or 8–12-week-old C57BL/6J mice.

To collect fully grown oocytes, ovaries were dissected in M2 medium supplemented with 2.5 µM milrinone (MilliporeSigma, M4659-10MG) to prevent spontaneous meiotic resumption. Preovulatory follicles were punctured to release fully grown oocytes. Cumulus-enclosed oocytes were collected for downstream analyses. Growing oocytes were mechanically released from secondary follicles using needles. Cumulus cells surrounding fully grown oocytes and granulosa cells surrounding growing oocytes were removed using hand-pulled glass capillaries. Oocytes were then cultured in M16 medium (MilliporeSigma, MR-010-D) with 2.5 μM milrinone before further experiments.

### In vitro culture of follicles and oocytes

In vitro culture of follicles was performed according to a previous report (Jaffe et al., 2011). Briefly, dissected follicles were transferred onto membrane inserts placed in 24-well plates pre-equilibrated with culture medium composed of MEM (STEMCELL Technologies, 36550) supplemented with 1× Penicillin-Streptomycin (Thermo Fisher, 15140122), 1× Insulin-Transferrin-Selenium (Thermo Fisher, 41400045), 5% (v/v) Fetal Bovine Serum (FBS; Thermo Fisher, 16000044), and 10 ng/mL follicle stimulating hormone (MilliporeSigma, F4021). Follicles were randomly assigned to treatment with either 5 μM nocodazole (MilliporeSigma, M1404-2MG) to depolymerize microtubules or DMSO (MilliporeSigma, D8418-50ML) as a vehicle control. The follicles were incubated in culture medium overnight, followed by fixation in 4% paraformaldehyde (PFA; Ted Pella, 18505) before subsequent immunostaining.

In vitro inhibitor treatments of oocytes isolated from secondary follicles were performed according to a previous report, with some modifications (Clift and Schuh, 2015). Briefly, oocytes were treated with 5 µM nocodazole in M16 medium for 1 or 2 h at 37°C, depending on the experimental objective, followed by thorough washing and incubation in M16 medium for 15 min at 37°C. Samples were subsequently fixed in 4% PFA for downstream analysis. To investigate the roles of the cytoskeleton in mitochondrial clustering, oocytes were stained with MitoTracker Deep Red (Thermo Fisher, M22426), and those exhibiting mitochondrial clustering were selected for subsequent experiments. Oocytes were randomly assigned to treatment with either 5 µM nocodazole to depolymerize microtubules, 1 µg/mL cytochalasin D (MilliporeSigma, C8273-1MG) to depolymerize F-actin, or DMSO as a vehicle control. The oocytes were treated for 2 h, followed by imaging to determine whether mitochondrial clustering was maintained. All experiments were performed in M16 medium supplemented with 2.5 μM milrinone to maintain meiotic arrest.

### MitoTracker, TMRM, and LysoTracker staining

MitoTracker, TMRM, and LysoTracker staining were performed according to the manufacturers’ protocols. Briefly, oocytes isolated from secondary follicles were incubated with 500 nM MitoTracker Deep Red (Thermo Fisher, M22426) for 30 min at 37°C, followed by thorough washing in fresh medium before imaging using a Leica Stellaris 8 DIVE confocal microscope. For TMRM (tetramethylrhodamine methyl ester) staining, oocytes were incubated with 1× Image-iT TMRM Reagent (Thermo Fisher, I34361) for 30 min at 37°C, followed by thorough washing and imaging using a Leica Stellaris 8 DIVE confocal microscope. For LysoTracker staining, oocytes were incubated with 50 nM LysoTracker Deep Red (Thermo Fisher, L12492) for 30 min at 37°C. For mitochondrial and lysosomal co-staining, MitoTracker Red CMXRos (Thermo Fisher, M7512) and LysoTracker Deep Red were used according to the manufacturers’ protocols. All experiments were performed in M16 medium supplemented with 2.5 μM milrinone to maintain meiotic arrest.

### Immunostaining

Whole-mount staining was performed as previously described (Li et al., 2017). Briefly, ovaries were fixed in 4% PFA at 4 °C overnight. Tissues were then sequentially incubated with primary antibodies for 2 days, washed for 1 day, and incubated with secondary antibodies for 2 days. Following immunostaining, samples were incubated in C_e_3D tissue-clearing buffer for one to two days prior to imaging with a Leica Stellaris 8 DIVE confocal microscope. Image data were analyzed using Fiji (ImageJ 1.54p) and Imaris 10.2.0.

Staining of isolated follicles and oocytes was performed as previously reported with modifications (Wang et al., 2025). Samples were fixed in a solution containing 100 mM HEPES (MilliporeSigma, H3375-25G), 50 mM EGTA (MilliporeSigma, E4378-25G), 10 mM MgSO (MilliporeSigma, MX0075-1), 2% PFA, and 0.5% Triton X-100 (MilliporeSigma, X100-500ML) in PBS at 37 °C for 10 min. After fixation, samples were washed overnight in 0.5% Triton X-100 in PBS, then blocked overnight at room temperature in blocking buffer composed of 3% (w/v) BSA, 5% (v/v) normal donkey serum (Jackson ImmunoResearch, 017-000-121), and 0.3% Triton X-100 in PBS. Samples were incubated with primary antibodies overnight at 4 °C and with secondary antibodies at room temperature for 1 h. Nuclei were counterstained with 1 µg/mL DAPI (MilliporeSigma, D9542-10MG) prior to imaging with the Leica Stellaris 8 DIVE confocal microscope. Higher laser power was used to visualize mitochondria. Image data were analyzed using Fiji (ImageJ 1.54p) and Imaris 10.2.0.

### Antibodies

anti-α-tubulin (Novus Biologicals, NB600-506, 1:1500, RRID:AB_10078394); anti-acetyl-α-tubulin (Lys40) (Cell Signaling Technology, 5335S, 1:1000, RRID:AB_10544694); anti-AURKA (Novus Biologicals, NBP1-51843SS, 1:100, RRID:AB_11018019); anti-CAMSAP1 (Proteintech, 83864-1-RR, 1:100, RRID:AB_3671450); anti-Calnexin (Abcam, ab22595, 1:100, RRID:AB_2069006); anti-CDK5RAP2 (MilliporeSigma, 06-1398, 1:100, RRID:AB_11203651); anti-CETN2 (Proteintech, 15877-1-AP, 1:100, RRID:AB_2077383); anti-CEP192 (Proteintech, 28700-1-AP, 1:100, RRID:AB_2918191); anti-CEP250/CNAP1 (Proteintech, 14498-1-AP, 1:100, RRID:AB_2076918); anti-DDX4 (Abcam, ab13840, 1:300, RRID:AB_443012); anti-DISC1 (Novus Biologicals, NB110-40773SS, 1:100, RRID:AB_1108155); anti-GM130 (BD Biosciences, 610822, 1:300, RRID:AB_398141); anti-HOOK3 (Proteintech, 15457-1-AP, 1:100, RRID:AB_2120828); anti-Lamin A/C (Abcam, ab238303, 1:300, RRID:AB_3722722); anti-Ninein (MilliporeSigma, ABN1720, 1:100); Phalloidin (Thermo Fisher, A12379, 1:400); anti-PCM1 (Proteintech, 19856-1-AP, 1:100, RRID:AB_2878616); anti-Pericentrin/PCNT (Abcam, ab4448, 1:300, RRID:AB_304461; BD Biosciences, 611814, 1:300, RRID:AB_399294); anti-PLK1 (Abclonal, A2548, 1:100, RRID:AB_2764437); anti-PLK4 (Proteintech, 28750-1-AP, 1:100, RRID:AB_2881206); anti-RAB9 (Thermo Fisher, MA5-31997, 1:100, RRID:AB_2809291); anti-TFAM (Proteintech, 22586-1-AP, 1:100, RRID:AB_11182588); anti-TOMM20 (Abcam, ab186735, 1:100, RRID:AB_2889972); anti-γ-tubulin (Abcam, ab11317, 1:300, RRID:AB_297921; MilliporeSigma, T3559, 1:300, RRID:AB_477575); Alexa Fluor 488 donkey anti-rat IgG (Thermo Fisher, A-21208, 1:600, RRID:AB_141709); Alexa Fluor 568 donkey anti-mouse IgG (Thermo Fisher, A10037, 1:600, RRID:AB_11180865); Alexa Fluor 647 donkey anti-mouse IgG (Thermo Fisher, A-31571, 1:600, RRID:AB_162542); Alexa Fluor 488 donkey anti-rabbit IgG (Thermo Fisher, A-21206, 1:600, RRID:AB_2535792); Alexa Fluor 568 donkey anti-rabbit IgG (Thermo Fisher, A10042, 1:600, RRID:AB_2534017); Alexa Fluor 647 donkey anti-rabbit IgG (Thermo Fisher, A-31573, 1:600, RRID:AB_2536183).

### Electron microscopy

Electron microscopy was performed as previously described (Yin and Spradling, 2025). Briefly, following initial fixation, ovaries were sequentially washed with 50 mM glycine (Millipore, G8898-500G) in 0.1 M PIPES buffer and with 0.1 M PIPES buffer (Electron Microscopy Sciences, 10010-67-0). Then tissue pieces were post-fixed in 1% osmium tetroxide (Electron Microscopy Sciences, 19190) and 1.5% potassium ferrocyanide (Electron Microscopy Sciences, 26604-01) for 1 h, washed in distilled water, and stained en bloc with 1% (w/v) uranyl acetate (Polysciences, 21447-25) for 1 h.

Samples were dehydrated through a graded ethanol series (30%, 50%, 75%, 85%, 95%, and 100%) followed by two exchanges in 100% acetone (Electron Microscopy Sciences, 10012). Ovaries were then infiltrated and embedded in EMbed 812 epoxy resin (Electron Microscopy Sciences, 14900) according to the manufacturer’s instructions. Ultrathin sections approximately 70 nm thick were cut using a Leica EM UC7 ultramicrotome and imaged with Hitachi HT7800/7600 transmission electron microscope operated under 80 kV. Images were acquired with an AMT NanoSprint 12 camera.

### Statistical analysis and study design

Quantitative measurements are presented as the mean ± SD of biological replicates. Categorical data are presented as the number and percentage of individual oocytes in each category. Statistical analyses were performed using GraphPad Prism 11 (GraphPad Software). Comparisons between two groups were performed using a two-tailed Student’s *t* test, or Welch’s *t* test, as appropriate. Comparisons among three or more groups were performed using one-way ANOVA, followed by Tukey’s multiple comparisons test. The chi-square test or Fisher’s exact test was used to analyze the association between categorical variables. The specific statistical test used for each analysis is indicated in the corresponding figure legend. Differences were considered statistically significant at *P* < 0.05. Statistical significance is indicated as follows: *P* < 0.05 (*), *P* < 0.01 (**), *P* < 0.001 (***), and *P* < 0.0001 (****).

No statistical methods were used to predetermine sample sizes. Sample sizes were determined based on prior experience and preliminary experiments. All experiments were performed at least twice. The authors were not blinded to the experimental design, conduct, or analysis. The numbers of biological replicates for each experiment are provided in the corresponding figure legends.

## ACKNOWLEDGEMENTS

We thank Dr. Mahmud Siddiqi for assistance with optical microscopy and image analysis, and Dr. Ru-Ching Hsia, Barbara J. Smith, Dr. Wanbao Niu, and Mike Sepanski for assistance with electron microscopy. We also thank the support staff of the Department of Embryology, Carnegie Institution for Science, and the Johns Hopkins University, as well as the current and former members of the Spradling laboratory, for their assistance and helpful discussions.

## Author contributions

Q.Y. and A.C.S. conceived the study, designed the experiments, analyzed the data, interpreted the results, and wrote the manuscript. All experiments were performed by Q.Y.

## Supplemental Figures

Figure S1. MitoTracker, TMRM, and LysoTracker staining and Golgi organization in mouse oocytes from secondary follicles. (A) Representative MitoTracker staining showing central enrichment of mitochondria in oocytes of secondary follicles. Scale bar, 20 μm. (B) Representative TMRM and LysoTracker staining showing central mitochondrial enrichment and modest enrichment of LysoTracker signal in oocytes of secondary follicles. Scale bar, 20 μm. (C) Immunostaining for GM130 and F-actin in an oocyte from a secondary follicle with a centrally located nucleus. DAPI (OE), high laser power and long exposure. Scale bar, 20 μm.

Figure S2. Changes in Bb organization in oocytes of primordial follicles. (A) Electron micrographs of primordial Bbs in the oocytes of primordial follicles. Examples are shown in which oocyte mitochondria in the Golgi-containing section are enriched around the Golgi apparatus (enriched), distributed relatively evenly throughout the cytoplasm (dispersed), or depleted from the region surrounding the Golgi apparatus (reversed). The proportions of sections with the indicated distributions are shown at right. The number above the bars indicates the number of primordial follicles analyzed. Red rectangles indicate regions enlarged below. Scale bars: top, 10 μm; bottom, 2 μm. (B) Representative immunostaining for GM130 in oocytes of primordial follicles from mice of different ages: P7 (postnatal day 7), 2 wk (2-week-old), 2 mo (2-month-old), and 4 mo (4-month-old). The GM130 channel alone is shown below, indicated by arrows. Scale bars, 20 μm. (C) Statistical analysis of the GM130 cluster in oocytes of primordial follicles during mouse development. Sample sizes (n) are indicated at the top of the panel. *P* value was determined using the Fisher’s exact test. ns, not significant. (D) Electron micrographs of centrioles in the oocytes of primordial follicles from P7 and P19 mice. Scale bars, 500 nm.

Figure S3. Microtubule acetylation in oocytes during primary follicle development. Representative images of three primary follicles at successive developmental stages showing changes in acetylated α-tubulin organization. Primary, primary follicle. Primary 1, with approximately 20 granulosa cells in the largest optical section, exhibits a centrally positioned nucleus and weak acetylated α-tubulin staining in the oocyte. Primary 2, with approximately 25 granulosa cells in the largest optical section, exhibits an eccentric nucleus and increased acetylated α-tubulin staining in the oocyte. Primary 3, with more than 30 granulosa cells in the largest optical section, exhibits an eccentric nucleus and the strongest acetylated α-tubulin staining in the oocyte. Dashed lines outline the nuclei. Arrows indicate MTOCs. Scale bars, 20 μm.

Figure S4. Golgi apparatus dynamics in oocytes during follicle development. Representative images showing the reorganization of the Golgi apparatus in an oocyte during follicle development. The Golgi apparatus forms a continuous Golgi ring in the oocyte of a primordial follicle, fragments while remaining enriched in the oocyte of a primary follicle, and becomes dispersed in the oocyte of a secondary follicle along with microtubule dynamics. Primordial, primordial follicle; Transitional, transitional follicle; Primary, primary follicle; Secondary, secondary follicle. Scale bars, 20 μm.

Figure S5. Inhibitor treatments of oocytes from secondary follicles. (A–C) Representative images showing that disruption of microtubules by nocodazole, but not disruption of F-actin by cytochalasin D, disperses the central mitochondrial cluster. n = 56 (DMSO), 29 (nocodazole), and 30 (cytochalasin D) oocytes, pooled from four independent experiments. *P* values were determined using a chi-square test of independence. Dashed lines outline the nuclei. Scale bars, 20 μm. (D) Representative images of oocytes from secondary follicles following 1 h of nocodazole treatment and 15 min of recovery. Arrows indicate MTOCs, which are enlarged in the insets together with the emanating microtubules. Scale bars: 20 μm; insets, 5 μm. (E) Statistical analysis of MTOC-associated microtubules per section. *P* value was determined using an unpaired two-tailed Welch’s *t* test. Sample sizes are indicated above the bars.

Figure S6. Three-dimensional reconstruction of follicles and effect of nocodazole treatment. (A) Left: Representative images of a secondary follicle with an eccentric nucleus and centrally localized MTOCs, and the corresponding three-dimensional reconstruction. Right: Representative images of a secondary follicle with a central nucleus and the corresponding three-dimensional reconstruction. Arrows indicate MTOCs. Secondary w/ eccentric, secondary follicle containing an oocyte with an eccentric nucleus; Secondary w/ central, secondary follicle containing an oocyte with a centrally localized nucleus. Scale bars, 20 μm. (B) Representative images of follicles following prolonged nocodazole treatment in vitro. Acetylated microtubules were largely depolymerized, accompanied by nuclear centralization and cortical redistribution of MTOCs. Dashed lines outline the oocytes. Arrows indicate MTOCs. Scale bar, 20 μm. (C) Statistical analysis of nuclear centralization in oocytes of follicles cultured in vitro with DMSO or nocodazole. Sample sizes are indicated above the bars. *P* values were determined using Fisher’s exact test. (D) Statistical analysis of MTOC location in oocytes of follicles cultured in vitro with DMSO or nocodazole. *P* values were determined using Fisher’s exact test. Sample sizes are indicated above the bars.

Figure S7. Fluorescence staining of MTOC-associated proteins. Representative immunofluorescence staining of MTOC-associated proteins, including AURKA (NSN = 19; SN = 15), CAMSAP1 (NSN = 22; SN = 3), CETN2 (NSN = 23; SN = 5), CEP192 (NSN = 16; SN = 4), CEP250 (NSN = 6; SN = 3), CDK5RAP2 (NSN = 16; SN = 6), DISC1 (NSN = 16; SN = 6), HOOK3 (NSN = 20; SN = 2), PLK1 (NSN = 21; SN = 6), and PLK4 (NSN = 20; SN = 3). All listed proteins were detected in the samples examined. Scale bars, 20 μm.

