## Supplemental Figures S1-S7 for "The mouse oocyte Balbiani body expands in growing follicles and influences nuclear position"

Figure. S1

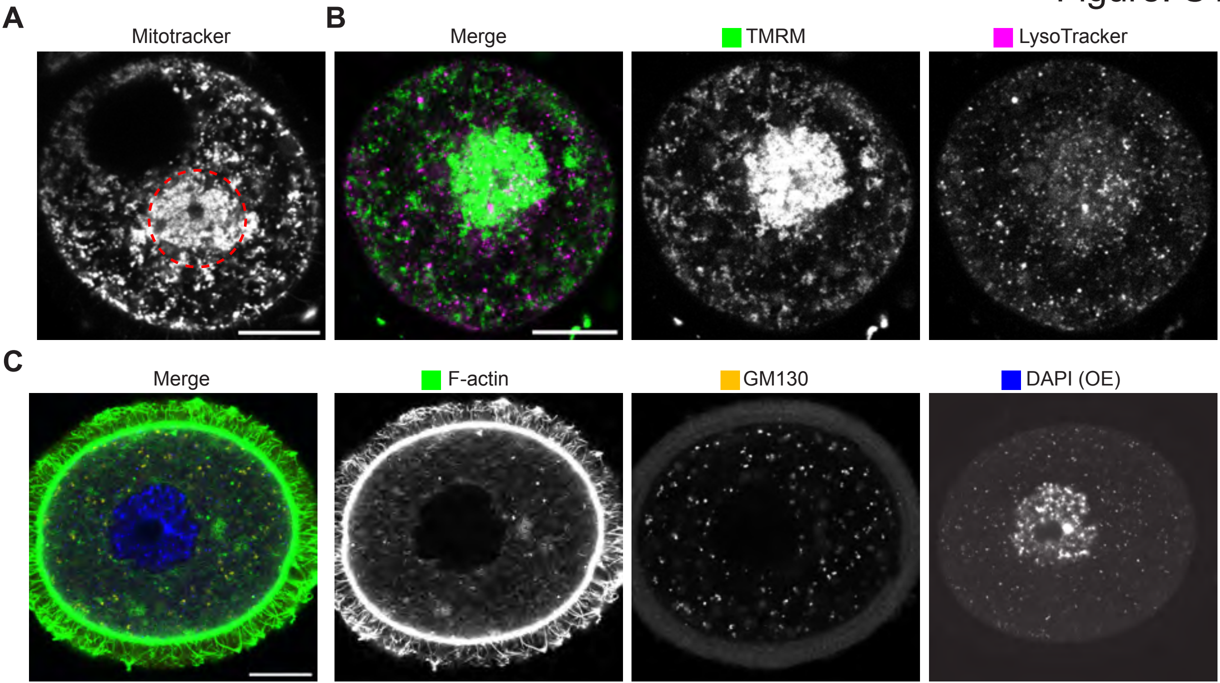
A
Mitotracker
B
Merge
TMRM
LysoTracker
C
Merge
F-actin
GM130
DAPI (OE)

Figure. S2

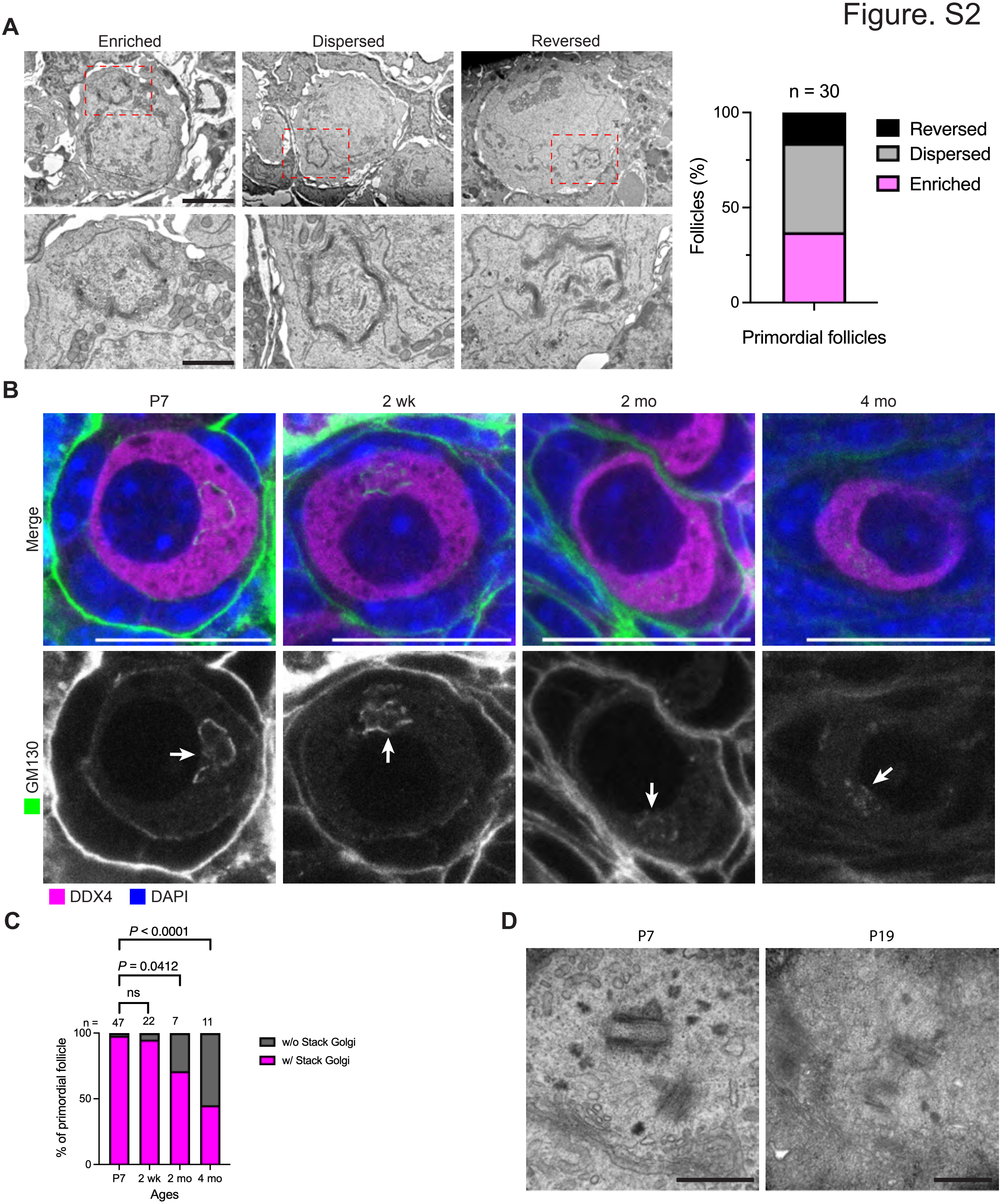
A
Enriched
Dispersed
Reversed
n = 30
100
50
0
Follicles (%)
Reversed
Dispersed
Enriched
Primordial follicles
B
P7
2 wk
2 mo
4 mo
Merge
GM130
DDX4
DAPI
C
P < 0.0001
P = 0.0412
ns
n = 47
22
7
11
100
50
0
% of primordial follicle
w/o Stack Golgi
w/ Stack Golgi
P7
2 wk
2 mo
4 mo
Ages
D
P7
P19

### Figure. S3

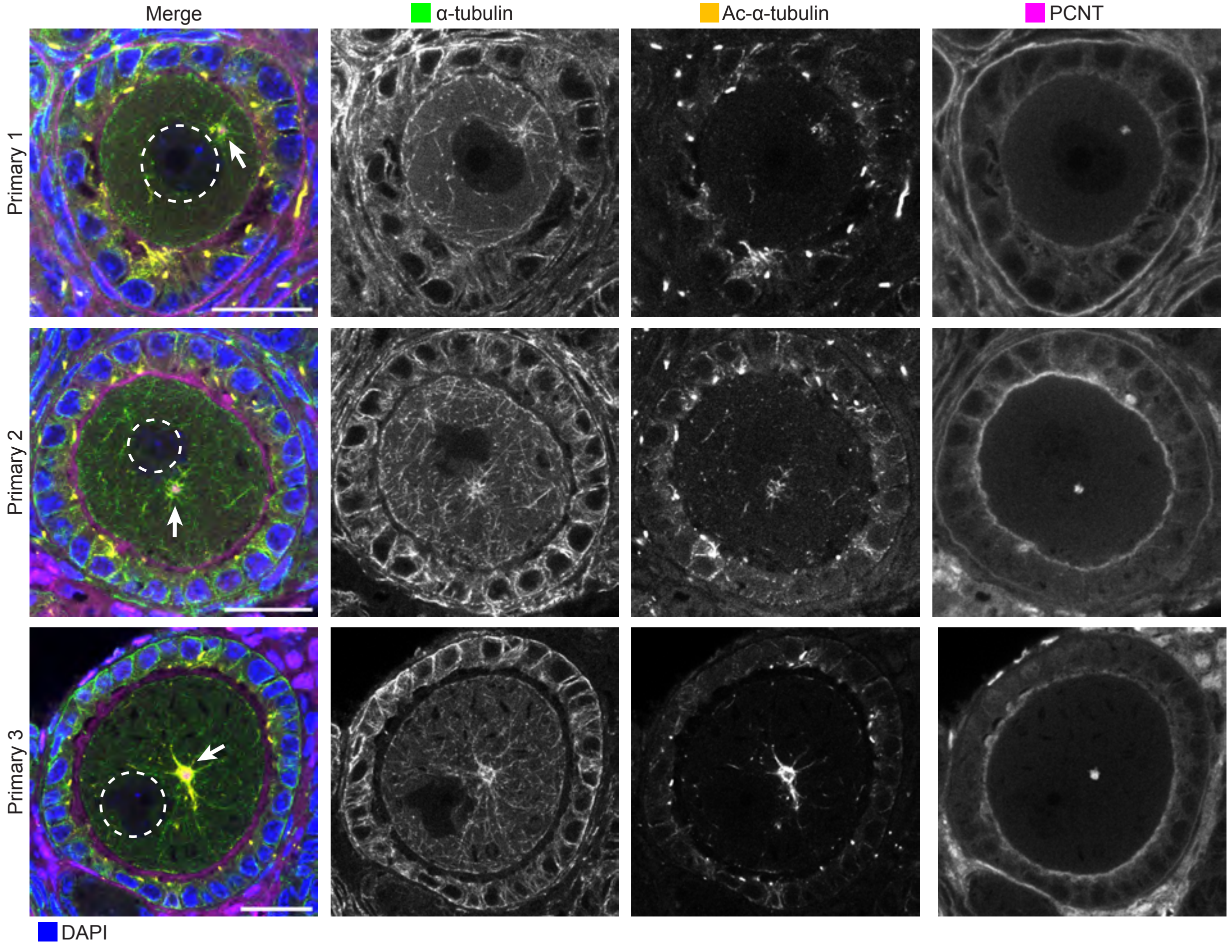
Merge
α-tubulin
Ac-α-tubulin
PCNT
Primary 1
Primary 2
Primary 3
DAPI

### Figure. S4

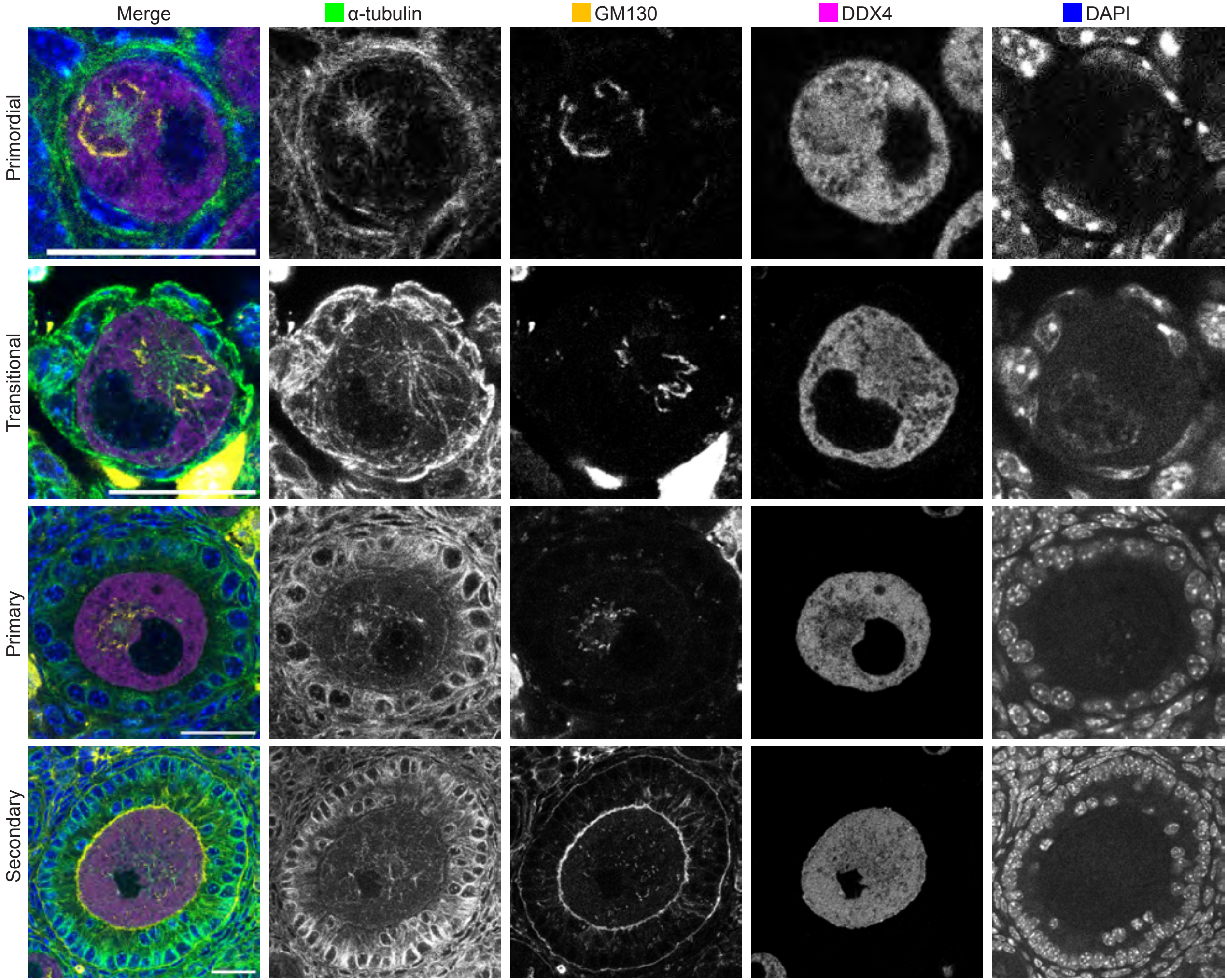
Merge
α-tubulin
GM130
DDX4
DAPI
Primordial
Transitional
Primary
Secondary

### Figure. S5

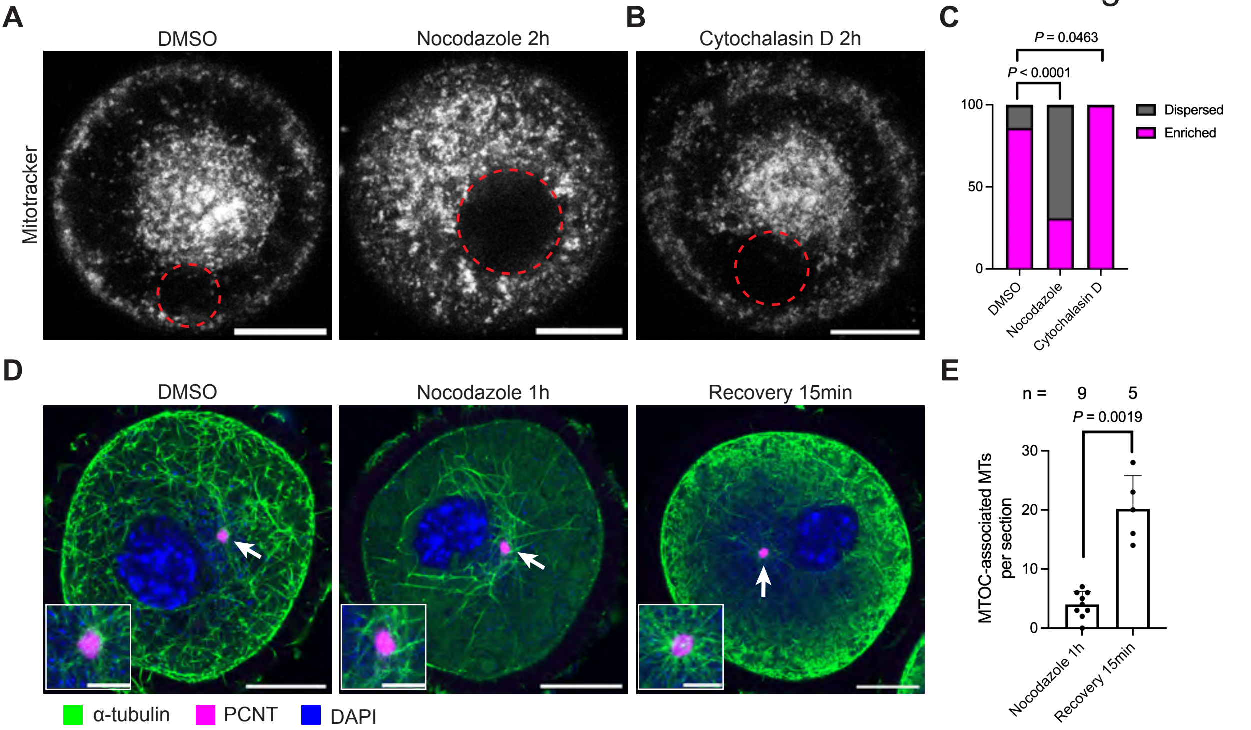
A
B
C
D
E
DMSO
Nocodazole 2h
Cytochalasin D 2h
Mitotracker
P = 0.0463
P < 0.0001
100
50
0
Dispersed
Enriched
DMSO
Nocodazole
Cytochalasin D
Nocodazole 1h
Recovery 15min
n = 9 5
P = 0.0019
30
20
10
0
MTOC-associated MTs per section
α-tubulin
PCNT
DAPI

### Figure. S6

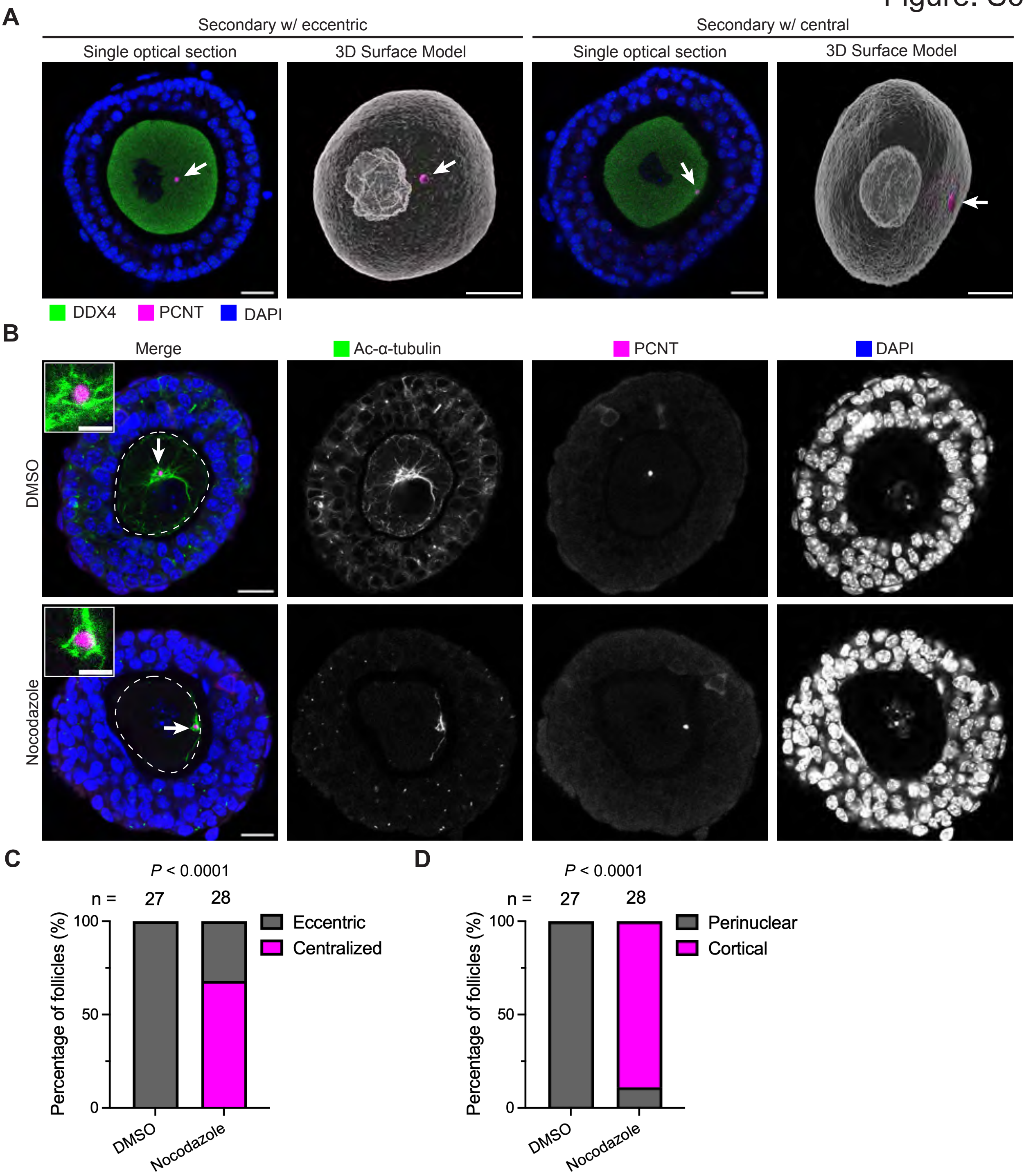
A
Secondary w/ eccentric
Secondary w/ central
Single optical section
3D Surface Model
Single optical section
3D Surface Model
DDX4
PCNT
DAPI
B
Merge
Ac-α-tubulin
PCNT
DAPI
DMSO
Nocodazole
C
P < 0.0001
n = 27 28
Percentage of follicles (%)
100
50
0
Eccentric
Centralized
DMSO
Nocodazole
D
P < 0.0001
n = 27 28
Percentage of follicles (%)
100
50
0
Perinuclear
Cortical
DMSO
Nocodazole

### Figure. S7

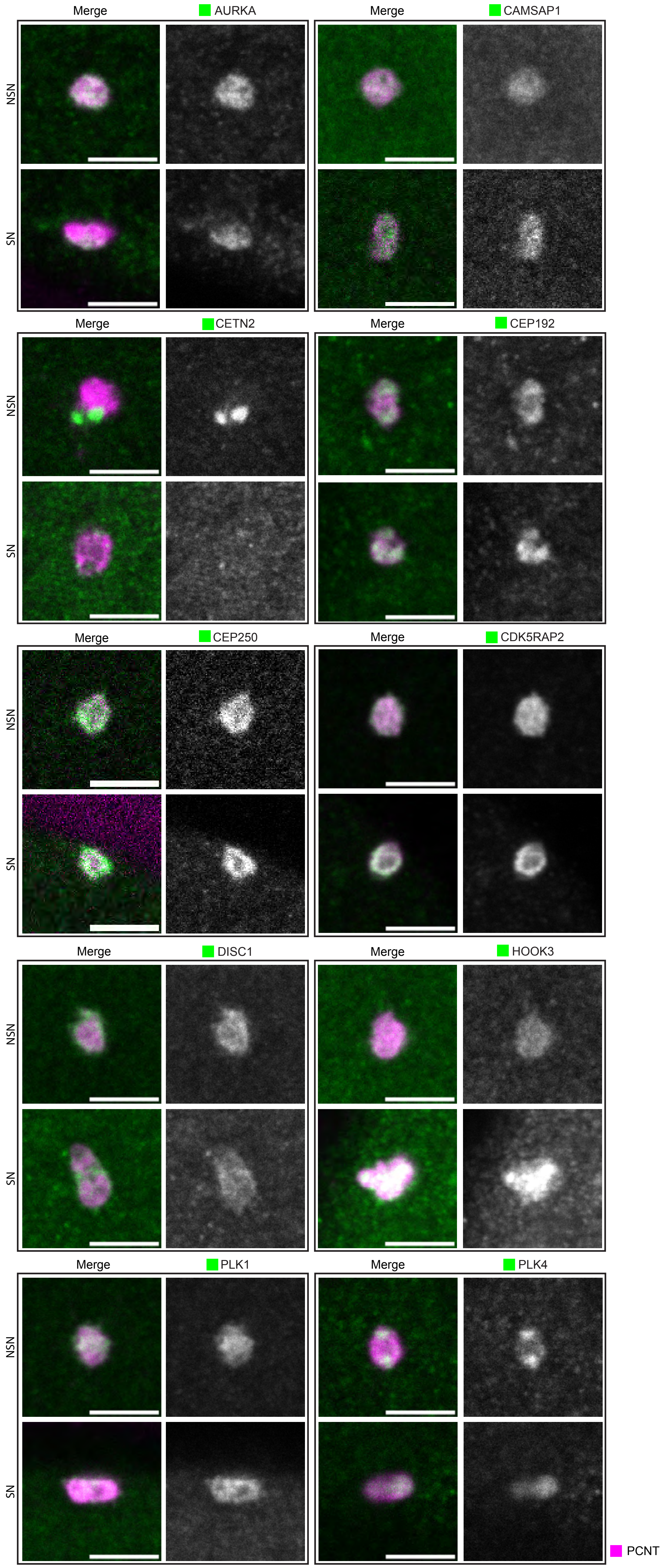
Merge
AURKA
Merge
CAMSAP1
NSN
SN
Merge
CETN2
Merge
CEP192
Merge
CEP250
Merge
CDK5RAP2
Merge
DISC1
Merge
HOOK3
Merge
PLK1
Merge
PLK4
PCNT
